# PlantTV: a versatile user-friendly software to explore RNA-seq data across plant species

**DOI:** 10.64898/2026.11.12.765996

**Authors:** Mila Pertuy, Barbara De Coninck, Lennard van Buren

## Abstract

**Background:** Bulk RNA sequencing (RNA-seq) has become a central tool in plant biology for studying genome-wide transcriptional responses. However, interpretation of differential expression results remains challenging, particularly for non-model species where functional annotation resources are fragmented, heterogeneous, or incomplete. Existing user-friendly RNA-seq analysis platforms often rely on predefined annotation databases, limiting their applicability across the diversity of plant species.

**Results:** We developed Plant Transcriptomics Viewer (PlantTV), a free web-based platform for interactive exploration of plant RNA-seq datasets. PlantTV integrates differential expression analysis, expression visualization, functional enrichment analysis, and co-expression network exploration within a single graphical interface. A central feature is the Annotation Builder, which allows users to generate reusable annotation tables from diverse resources, including GFF/GTF files, EggNOG annotations, BLAST results, and custom tabular datasets. Using case studies in *Arabidopsis thaliana, Malus domestica*, and *Eucalyptus melliodora*, we demonstrate that PlantTV supports transcriptome analysis across a range of annotation scenarios, from well-annotated reference genomes to de novo transcriptome assemblies.

**Conclusions:** PlantTV provides an interactive and species-independent environment for plant transcriptome analysis. By combining RNA-seq exploration tools with flexible support for custom annotation resources, PlantTV facilitates biological interpretation of transcriptomic datasets in both model and non-model plant species.

## Background

In the past decade, bulk RNA-sequencing (RNA-seq) has become a cornerstone of plant molecular biology, enabling genome-wide measurements of gene expression in a single experiment. Its widespread adoption is driven by several advantages, including technical simplicity, low cost, modest requirements for biological input material, and the broad applicability of poly(A)-based library preparation across species (Upton et al. 2023; Stark et al. 2019). In parallel, the growing availability and improving quality of reference genomes further facilitates RNA-seq analysis across a wide range of plant systems (Sun et al. 2022; Michael and VanBuren 2020).

While generating an RNA-seq dataset is nowadays relatively straightforward, data interpretation often remains a bioinformatics challenge. Following read alignment and quantification through standardized analysis pipelines, differential expression analysis is commonly used to identify genes whose expression changes significantly between conditions (Love et al. 2014). However, a typical experiment may yield hundreds to thousands of differentially expressed genes (DEGs) across multiple treatments, genotypes or time points. Extracting biological insight from these large DEG sets often requires a range of complementary downstream approaches and visualizations, including volcano plots, heatmaps, Venn diagrams and overrepresentation analysis (McDermaid et al. 2019; Conesa et al. 2016).

The value of differential expression analysis lies not only in identifying DEGs, but also in understanding their biological function. Functional genome annotations, where gene identity is linked to predicted gene function, provide the context necessary to uncover the biological processes that underly transcriptomic changes. Consequently, the biological interpretation of transcriptomic datasets is depends on the availability, quality and accessibility of functional annotation resources.

This dependence poses a particular challenge in plant biology, where researchers work across thousands of species that differ greatly in the availability and quality of genomic resources. While model species often benefit from well-maintained annotations available through centralized repositories such as NCBI and Ensembl, many non-model species rely on alternative resources, including community databases, de novo transcriptome assemblies, orthology-based annotation pipelines, and project-specific annotation efforts. Functional information may therefore be available in a variety of formats, including GFF annotations, Gene Ontology assignments, EggNOG predictions, Trinotate outputs and custom annotation tables generated within research projects.

The diversity of annotation resources in plant biology is not only a consequence of incomplete database coverage, but also reflects the inherent complexity of plant genomes. Large genome sizes, widespread whole-genome duplication events, extensive gene family expansions, abundant transposable elements and variable ploidy levels complicate both genome assembly and functional annotation (Michael 2026; Vuruputoor et al. 2023). As a result, multiple annotation resources frequently coexist for a single species, each differing in scope, methodology, or intended application. Researchers may therefore prefer different resources dependent on the scope of their project or their species of interest, or combine complementary annotation resources to obtain a more complete functional understanding.

Several user-friendly platforms, such as iDEP (Ge et al. 2018) and DEBrowser (Kucukural et al. 2019) provide graphical user interfaces (GUIs) to support differential expression analysis of RNA-seq data and downstream interpretation. Despite their utility, these platforms are often built around predefined organisms and annotation resources, which limits their flexibility when alternative or user-generated annotations are required as is the case for many plant species. Although workflow management systems such as Galaxy (The Galaxy Community 2026) offer greater flexibility in designing analyses, they introduce unnecessary complexity when the primary objective is the exploration of differential expression results. Moreover, these workflow managers are not designed to let users easily navigate between complementary analyses, and thus often require researchers to manually integrate the results from different plots and visualizations.

While downloadable analysis packages or custom-built computational workflows can overcome some of these limitations, they typically produce static outputs that are not suited for exploratory data analysis. Researchers often need to query specific genes, gene families, or pathways within their dataset. In addition, transcriptome analysis is rarely a linear process, as new findings and hypotheses frequently drive researchers to revisit existing datasets. To accommodate this iterative, hypothesis-driven data exploration, easy and intuitive user interaction with their dataset is required.

The aforementioned challenges highlight the need in plant science for an easy-to-use transcriptome data analysis platform that combines interactive data exploration with flexible integration of diverse annotation resources. To address this need, we developed Plant Transcriptomics Viewer, or PlantTV, a free web-based platform for plant RNA-seq data analysis. PlantTV provides a centralized and interactive environment for the exploration of differential expression results and supports the custom integration of a range of genome annotations. By enabling flexible and annotation-aware transcriptome analyses, PlantTV aims to facilitate biological discovery beyond model plant species.

### Implementation

#### Software architecture

We developed PlantTV in Python using the open-source Panel framework, which provides support for interactive web applications through integrated widgets, layouts, and data visualization components. We designed a modular architecture in which different analyses and visualizations are implemented as independent components, and represented in the GUI across different tabs. This modular design facilitates future extension of the platform while minimizing dependencies between analysis modules. Interactive visualizations are implemented using HoloViews and Bokeh, enabling dynamic user interactions, including gene and pathway selection, sample filtering, and retrieval of additional information through hover functionality. We used Claude AI for iterative GUI development, code debugging, and improving the robustness of user experience.

PlantTV is built around two principal data objects: *studies* and *annotations*. This separation allows RNA-seq datasets and gene annotation resources to be managed independently, thereby enabling the reuse of species annotation information across multiple studies.

#### Study structure and differential expression analysis

A study in PlantTV contains all experiment-specific information required for downstream analyses and consists of four input files: (1) a *sample sheet* describing sample identities, experimental groups and any additional metadata, (2) a gene-level *count matrix*, (3) a *contrasts file* defining pairwise comparisons between sample groups, and (4) a *differential expression results* file generated with DESeq2. Contrasts are specified as *groupA*.*vs*.*groupB* comparisons using sample groups defined in the sample sheet.

To simplify data preparation, PlantTV provides a built-in *Study Builder* that guides users through the creation of a study object. All input files are automatically validated for compatibility and formatting, and PlantTV integrates pyDESeq2 to perform differential expression analyses directly from uploaded count matrices and contrast definitions.

#### Flexible annotation framework

PlantTV supports the use of a wide variety of annotation resources. Annotation information is stored separately from study data and linked to species through metadata, allowing annotation resources to be re-used across datasets. Annotation tables can contain a variety of gene-associated information, including gene symbols, functional descriptions, Gene Ontology (GO) terms, KEGG pathway annotations, and other user-defined metadata.

To help researchers generate structured annotation information from diverse sources, PlantTV includes an integrated *Annotation Builder* module. The Annotation Builder consolidates heterogeneous gene information which is often distributed across multiple resources, including genome annotation files, functional annotation pipelines, homology searches and manually curated datasets, into a single, standardized annotation table. Per result, users can combine gene symbols, functional descriptions, pathway annotations, and other custom metadata into a unified gene-centric annotation resource that is subsequently available throughout all PlantTV’s analysis modules.

The Annotation Builder supports multiple commonly used input formats, including genome annotation files (GTF and GFF/GFF3), BLASTp homology searches (outfmt6) (Altschul 1997) and eggNOG-mapper annotation outputs (.emapper.annotations) (Cantalapiedra et al. 2021). In addition, support of generic tabular formats (.csv, .tsv, .xlsx) also allows incorporation of tabular output files generated by genome assembly and annotation workflows, including Trinity and Trinotate (Haas et al. 2013; Bryant et al. 2017). Users may additionally augment annotation tables with custom attributes obtained from external databases or manual curation efforts.

Annotation completeness varies considerably among species and research projects. PlantTV is therefore designed to operate across different levels of annotation completeness. Core analytical functions remain available in the absence of extensive annotation information, while additional functionalities, such as functional enrichment analysis and enhanced gene querying, only become available when the relevant information is present in the selected annotation table.

#### Deployment, data management and accessibility

PlantTV supports analysis of private datasets in secure, user-authenticated environments as well as public datasets accessible to all visitors. User-uploaded studies can be made public upon request to promote dissemination and reuse of transcriptomic datasets. The platform is deployed as a containerized web application using Docker on university servers and is freely accessible through a web interface without requiring local software installation or programming expertise. Comprehensive documentation and user guides can be found online.

## Results & Discussion

PlantTV is a platform that integrates transcriptome analysis workflows and flexible annotation resources within a user-friendly graphical interface. The tool enables exploration of differential expression results and functional annotations without requiring programming expertise (Fig. 1).

**Fig. 1.**
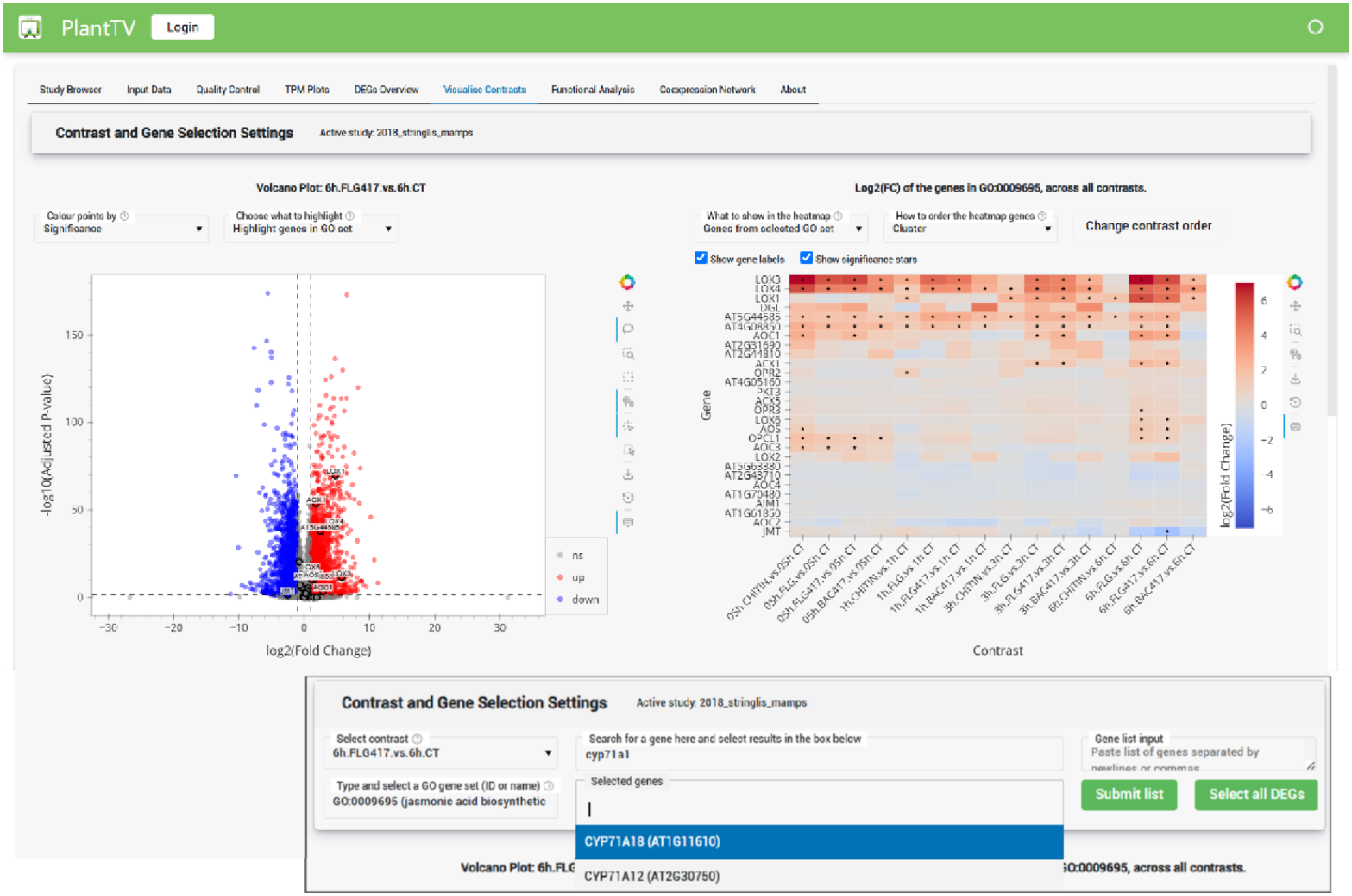
Example snapshot of PlantTV’s graphical user interface. PlantTV provides a comprehensive set of functionalities relevant for differential gene expression analysis. Different analyses and visualizations are split across multiple tabs. A study is first loaded in the Study Browser, accompanied by a user-chosen annotation table, after which all other tabs become populated. From left to right: Input Data, Quality Control, TPM Plots, DEGs Overview, Visualise Contrasts, Functional Analysis and Coexpression Network. Each tab contains a different analysis or visualization, and the data can be interactively queried by the user. Depicted here is the Visualize Contrasts tab, where *A. thaliana* root transcriptome data (Stringlis et al. 2018) is loaded and the differential expression results for a single treatment and timepoint are plotted in a volcano plot and heatmap. User inspection of the GO-term “Jasmonic acid biosynthesis” (GO:0009695) highlights the associated genes in the volcano-plot (left) and displays expression profiles in these genes in the heatmap (right). Inset: example form for data querying. An interactive widget allows to search GO terms, gene symbols, or paste gene lists, after which corresponding genes are highlighted in the volcano and plotted in the heatmap.

### Comparison with other tools

While PlantTV incorporates many of the visualizations and analysis capabilities found in platforms such as DEBrowser and iDEP, its key innovation is support for user-defined annotation resources. Using the integrated Annotation Builder, researchers can construct annotation tables from multiple sources and combine them with additional custom gene information. This flexibility enables downstream analysis and querying for both model and non-model plant species, independent of the availability of predefined genome annotations in centralized databases. PlantTV further leverages the annotation information for advanced querying, allowing users to search genes, gene lists, and pathways using human-readable identifiers and immediately examine their expression patterns. This makes gene lookup intuitive, which is crucial for interactive data exploration. Another difference with existing tools is that PlantTV hosts both public and private datasets. Datasets can be made public upon requests, allowing researchers to share their data in an accessible way with the wide plant science community. A full comparison between PlantTV and other RNA-seq analysis tools can be found in Table 1.

**Table 1.**
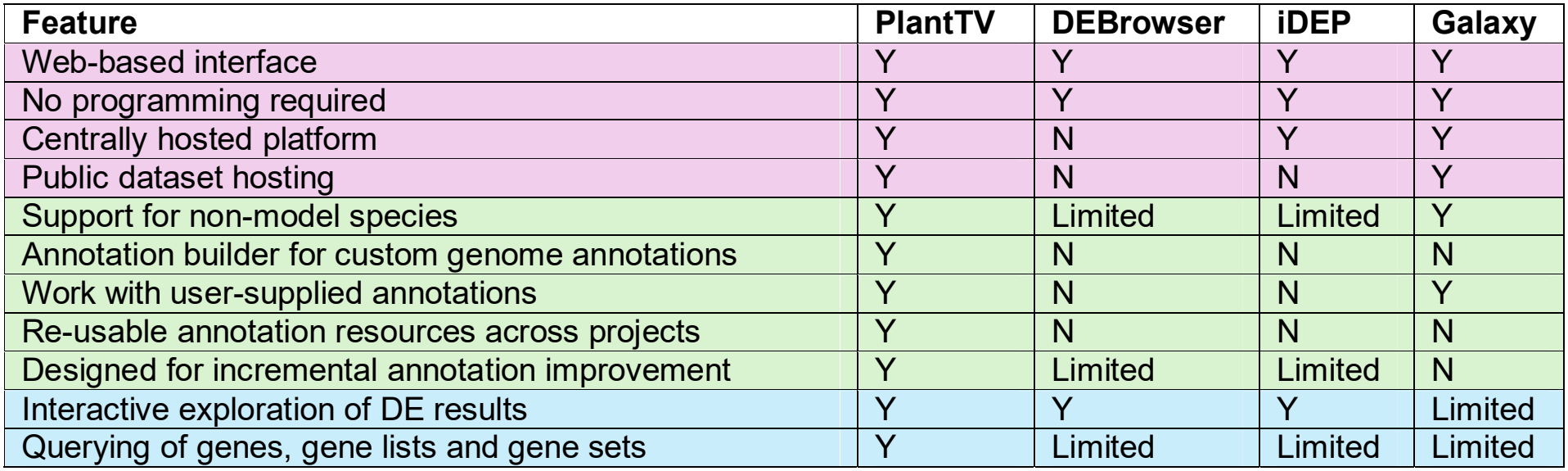
Comparison of PlantTV with other RNA-seq data analysis platforms. Features are grouped into three categories: hosting and accessibility (pink), support across plant species (green), and user interaction (blue).

### User workflow and analytical modules

The overall workflow in PlantTV consists of study selection, annotation loading, and dynamic exploration of transcriptomic patterns through its analysis modules (Fig. 2). Available datasets can be searched through the *Study Browser* using study metadata such as species, treatment and other keywords. Following data loading and pairing with an annotation table, users can access all study inputs, including sample metadata, count matrices, differential expression results and annotation table through the *Input Data* module. PlantTV further provides quality control visualizations in the *Quality Control* module, including principal component analysis (PCA) and chromosome- or contig-level read distribution plots, which together enable rapid assessment of sample relationships and potential technical issues.

**Fig. 2.**
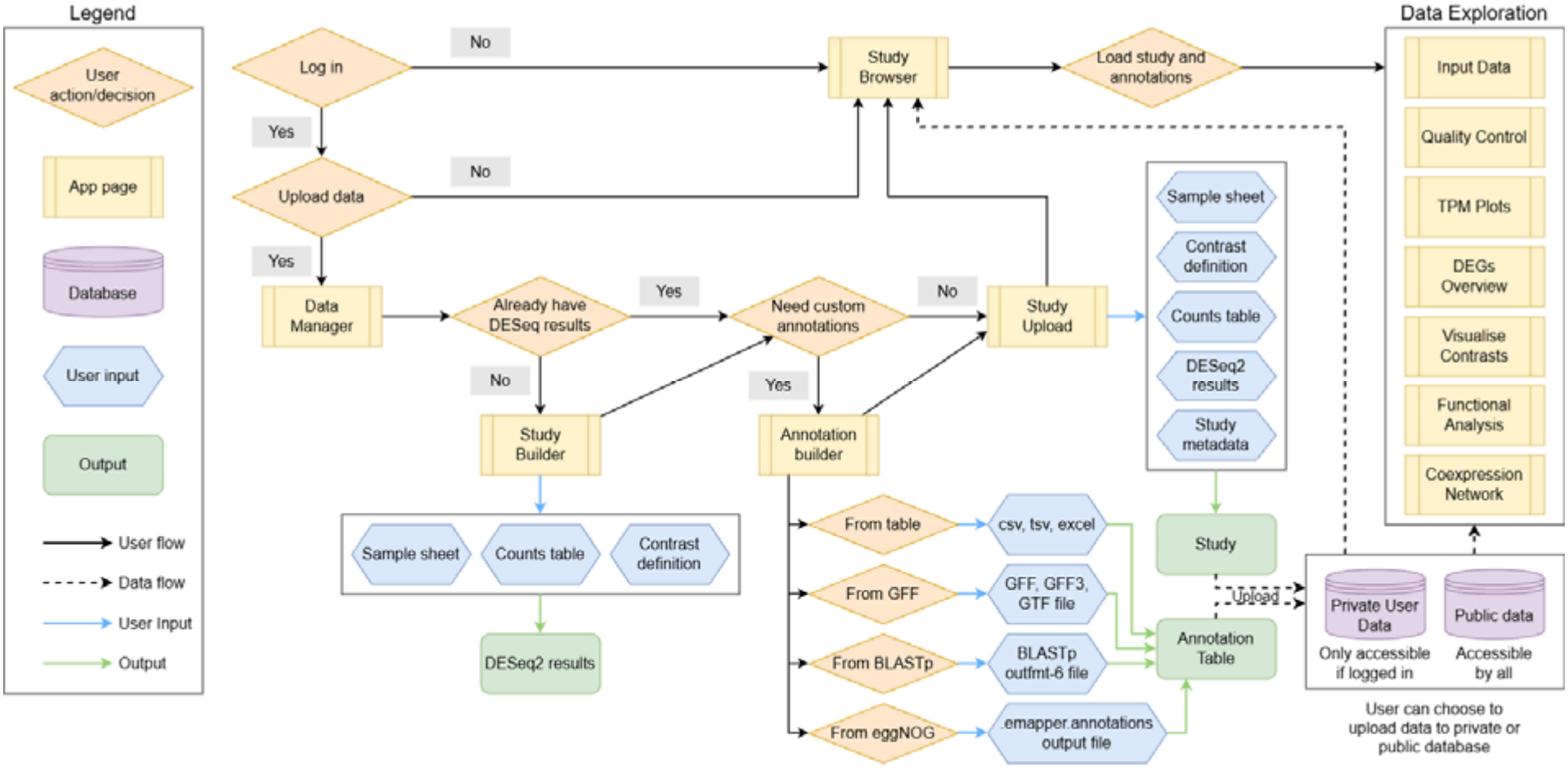
Schematic overview of user workflow. Without logging in, users can load public studies and annotations from the Study Browser and explore these studies using all PlantTV’s data exploration tabs. Upon login, users additionally get access to the Data Manager tab, which enables them to build studies and annotations, and upload these to their private repository in PlantTV.

Transcript abundance patterns can be explored in the *TPM Plots* module, which visualizes transcript levels for requested genes across samples, treatments, developmental stages or time series. The *DEGs Overview* module visualizes total numbers of up- and downregulated DEGs across contrasts and further supports overlap analysis using Venn diagrams and Upset plots, allowing rapid identification of shared and contrast-specific responses. One can inspect intersections to reveal constituent genes and their available functional information.

A central feature of PlantTV is the *Visualize Contrasts* module, which shows a volcano plot and heatmap side-by-side in a linked environment. Genes can be selected directly from the volcano plot, and their expression examined simultaneously in the heatmap across all relevant contrasts. In addition to manual selection, users may query individual genes, upload custom gene lists, or interrogate functional categories that are assembled from the annotation table. Several visualization schemes are available, including gene colouring based on the number of contrasts in which differential expression is observed which quickly reveals broadly responsive and condition-specific genes.

Functional interpretation of transcriptional responses is provided through the *Functional Analysis* module. Gene Ontology (GO) and KEGG enrichment analyses are performed on differentially expressed gene sets through the *GSEApy* implementation of Enrichr (Chen et al. 2013; Kuleshov et al. 2016; Fang et al. 2023), provided that gene set information is provided in the annotation table. Significantly enriched terms are visualized interactively and can be explored further to identify the underlying genes and their expression patterns.

Since annotation-based analyses are inherently constrained by the available functional knowledge, PlantTV additionally incorporates a *Co-expression Network* module to support exploratory analyses. For *Arabidopsis thaliana*, co-expression relationships derived from the ATTED-II database (Ath-r.c7-0) (Obayashi et al. 2022) are used to construct networks from differentially expressed genes. Nodes represent genes, while edges indicate co-expression relationships that exceed a user-defined threshold. These networks facilitate identification of transcriptionally coordinated gene clusters and may uncover functional associations that are not captured by traditional enrichment analyses.

After logging in, users get additional access to the *Data Manager* tab. In this tab, users can build studies and annotation tables, which they can then upload to PlantTV. This gives users the option to analyse their own data, of any plant species, in a private environment.

Collectively, these interconnected modules provide a comprehensive framework for transcriptome analysis, spanning quality assessment, expression profiling, differential expression analysis, functional interpretation, and co-expression analysis. By integrating these functionalities in a single interactive environment, combined with the added annotation support, PlantTV enables efficient and customizable exploration of plant RNA-seq data.

#### Use cases

We benchmark PlantTV and demonstrate its versatility through three use cases that represent common challenges in plant transcriptomics analysis across diverse species and annotation frameworks. These include reanalysis of an RNA-seq dataset from the well-characterized *Arabidopsis thaliana*, validation of transcriptome analysis using a custom annotation for *Malus domestica* assembled from community resources, and construction of a functional annotation table directly from EggNOG-mapper output. Together, these examples illustrate how PlantTV supports both model and non-model plant species and enables analysis regardless of whether annotations originate from curated reference databases, community-generated resources, or de novo functional annotation pipelines.

##### 1. Well-annotated genome available: *Arabidopsis thaliana*

To demonstrate PlantTV’s suite of analyses modules, we reanalysed a published transcriptomic dataset examining *Arabidopsis thaliana* root responses to beneficial and pathogenic microbial stimuli (Stringlis et al. 2018). As no publicly available gene count matrix was provided, we download raw sequencing reads from the Short Read Archive (BioProject ID: PRJNA412447) and aligned these to the *Arabidopsis* reference genome (TAIR10) using the nf-core RNA-seq pipeline (Ewels et al. 2020) to generate a count table. We then performed differential expression analysis using PlantTV’s built-in pyDESeq2 in the Study Builder module and uploaded the dataset to PlantTV. Both the processed study and the associated *A. thaliana* annotation are publicly accessible through the PlantTV platform.

Following study loading, we first verified that transcript abundance estimates (TPM values) reproduced the expression patterns reported in the original publication after root treatment with the flagellin epitope flg22 originating from the plant-pathogenic bacterium *Pseudomonas syringae* and beneficial bacterium *Pseudomonas simiae* WCS417, flg22^Pa^ and flg22^417^, respectively. Both flagellin epitopes were reported to sharply increase CYP71A12 levels after 3h and 6h treatment time, which is also captured in PlantTV’s TPM plot (Fig. 3a).

**Fig. 3.**
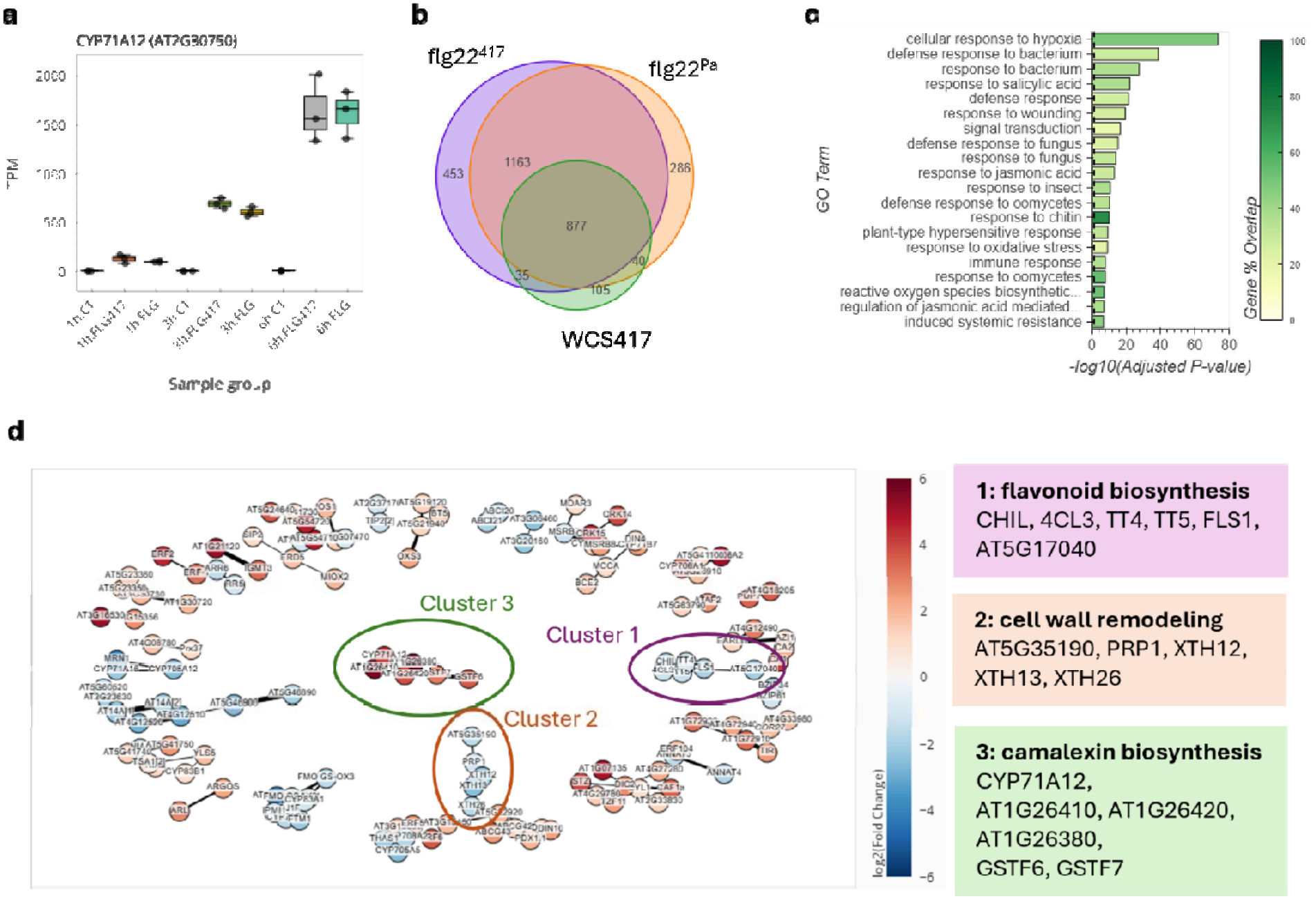
Representative PlantTV visualizations generated from a public *Arabidopsis thaliana* transcriptomic dataset. Data were derived from *A. thaliana* root transcriptome profiles (Stringlis et al. 2018). Panels shown are representative outputs generated by PlantTV. Some panels have been adjusted for visualization purposes. **(a)** TPM boxplots for CYP71A12 across a subset of selected experimental contrasts. Points represent individual replicates, boxes indicate the interquartile range, and the central line denotes the median. **(b)** Venn diagram showing the overlap of DEGs induced by flg22^417^, flg22^Pa^ and WCS416 after 1h of treatment time. **(c)** Gene Ontology enrichment analysis for upregulated genes after 1h treatment with flg22^417^. The 20 GO terms with the lowest adjusted P-values are shown. **(d)** Co-expression network for genes differentially expressed after 1h of treatment with flg22^417^. Nodes represent genes, edges represent co-expression relationships with z-score exceeding the user-defined threshold of 13. Node colour indicates log2 fold-change in gene expression. Three clusters are highlighted and discussed in the main text.

**Fig. 3.**
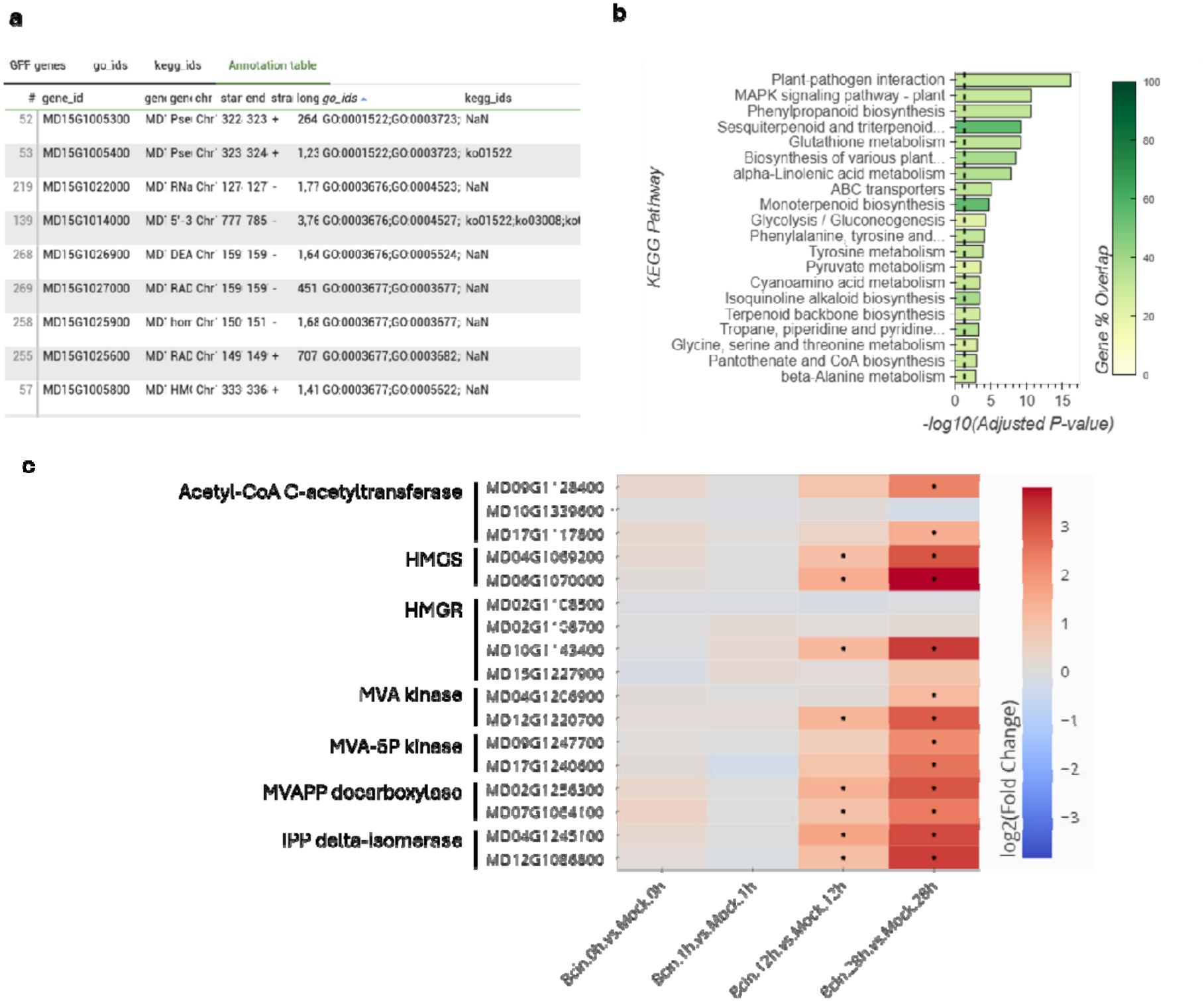
Analysis in PlantTV using a custom-generated annotation table. Different genomic resources were compiled in PlantTV’s annotation builder to support analysis of RNA-seq data of *Malus domestica* fruit infected with *Botrytis cinerea*. **(a)** Screenshot of the Annotation Builder showing the annotation table after automated parsing of the *M. domestica* GFF file and merging with InterProScan and KAAS output. **(b)** KEGG overrepresentation analysis at 28hpi performed with significantly upregulated genes. **(c)** Heatmap showing expression patterns of genes in the mevalonate biosynthesis pathway, a key upregulated pathway identified in the original publication. Significantly deregulated genes are highlighted with an asterisk. Gene identifiers and their functional families were obtained from the original publication and queried in PlantTV using the gene list lookup widget.

Then, we looked at overlap in differentially expressed genes. Per example, we inspected DEGs after 1h of treatment time. Both flagellin epitopes induced a similar gene expression response as also captured by PlantTV’s Venn diagram, with an overlap of over 2000 DEGs after 1h treatment time (Fig. 3b). Root colonization by WCS417 induced a similar, but weaker response according to the authors, which is also reflected in the overlapping yet lower number of 877 DEGs of WCS417-treated with both flg22-treated roots (Fig. 3b). Functional enrichment analysis within PlantTV confirmed a strong induction of stress-response pathways among genes upregulated after 1h of flg22 treatment (Fig. 3c). These findings further corroborate the biological conclusions reported in the original paper.

Finally, we applied the co-expression visualization module to identify functionally coherent gene clusters. Examination of DEGs following 1h of flg22^417^ treatment revealed multiple distinct co-expression clusters (Fig. 3d). To optimize network resolution, we varied the z-score threshold for co-expression and selected a threshold of 13, which produced a network containing multiple clearly separable clusters. As expected, genes within individual clusters exhibited coordinated expression patterns, indicating that the co-expression relationships in this study are consistent with those reported in the ATTED-II database. Several biologically meaningful modules could be readily identified. One cluster of genes of downregulated genes contained TT4, TT5, FLS1 and 4CL3, all of which are involved in flavonoid biosynthesis (Yonekura-Sakakibara et al. 2008). Another downregulated cluster contained genes associated with cell wall remodelling, including XTH12, XTH13, XTH26 and PRP1. These findings are consistent with the results of enrichment analysis (Fig. S1). In addition, we identified a strongly upregulated cluster containing CYP71A12, GSTF6, GSTF7 and several other genes that are all involved in or associated with camalexin biosynthesis, an important phytoalexin produced by *A. thaliana* (Millet et al. 2010; Su et al. 2011). Notably, camalexin biosynthesis was not among the top 20 enriched Gene Ontology terms (Fig. 3c), which illustrates the value of co-expression network visualization as a complementary perspective to functional enrichment analysis.

##### 2. Compile annotation file from multiple community resources: *Malus domestica*

To demonstrate PlantTV’s applicability to non-model species, we analysed an RNA-seq dataset investigating the molecular response of apple (*Malus domestica)* fruit to infection by the necrotrophic fungal pathogen *Botrytis cinerea* (Naets et al. 2022). The raw counts table was downloaded from NCBI Gene Expression Omnibus (GSE195841) and processed with PlantTV’s Study Builder. To inspect these data in PlantTV, we first obtained genomic resources for *M. domestica* from the Genome Database for Rosaceae (GDR) (Jung et al. 2019), consisting of three separate files: a GFF annotation file, a gene-to-Gene Ontology (GO) mapping generated using InterProScan (Jones et al. 2014), and a gene-to-KEGG mapping obtained through the KEGG Automated Annotation Server KAAS (Moriya et al. 2007). These resources were combined into a single annotation table and linked to the RNA-seq dataset in PlantTV to support downstream analysis (Fig. 4a).

Following the upload of study data and the generated annotation table, we explored the transcriptomic responses using PlantTV’s functional analysis tools. Because we built a detailed annotation table, gene information was available throughout PlantTV for various analyses. At 28 hours post-inoculation (hpi), KEGG pathway enrichment analysis revealed a clear induction of defence-associated pathways, including MAPK signalling and phenylpropanoid biosynthesis (Fig. 4b. These results are consistent with the biological conclusions reported in the original publication, demonstrating that PlantTV can rapidly reproduce key findings from transcriptomics studies in non-model species.

In addition to pathway-level analyses, PlantTV enabled direct inspection of individual genes and gene lists through the gene query widgets. To demonstrate this, we inspected the expression profile of genes in the mevalonate pathway, which were highlighted and experimentally validated by Naets et al. We obtained a list of gene identifiers from the original publication, and pasted this list into the gene lookup widget in the *Visualise Contrasts* tab (Fig. 1, inset). This readily revealed their significant upregulation especially at later timepoints during infection. Together, these analyses illustrate how PlantTV provides an intuitive framework for both functional interpretation and targeted gene-level exploration of transcriptomic datasets in species where annotation resources are not available in centralized databases.

##### 3. De novo transcriptome assembly: Eucalyptus

In species lacking community-supported genome annotations, transcriptome analyses often rely on de novo transcriptome assemblies. To demonstrate PlantTV’s applicability in this setting, we used transcriptomic resources from an *Eucalyptus* species, *Eucalyptus melliodora*, generated through de novo transcriptome assembly (Ferguson et al. 2024). We obtained EggNOG output files from the public repository Figshare (https://doi.org/10.6084/m9.figshare.24616131). EggNOG annotation outputs were successfully converted into PlantTV annotation tables containing functional descriptions, Gene Ontology (GO) terms, and KEGG pathway information (Fig. 5). After uploading this file to PlantTV’s repository, extended gene information is available across downstream analyses. This example illustrates how PlantTV can support transcriptome exploration in species for which annotation resources originate from de novo annotation pipelines rather than established community databases.

**Fig. 5.**
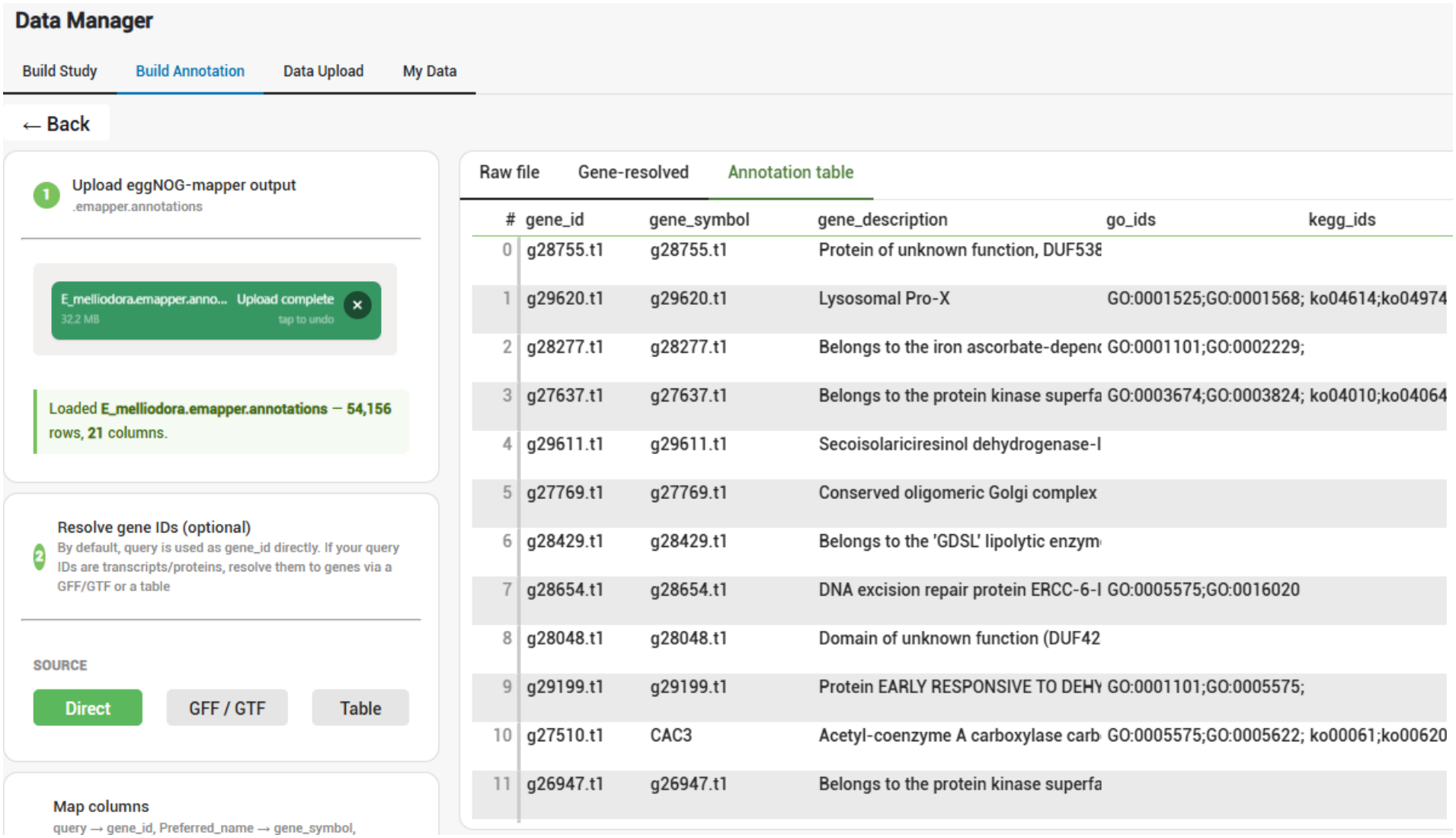
Screenshot of the Annotation Builder processing EggNOG mapper output. After loading the EggNOG output of *E. melliodora* into PlantTV’s Annotation Builder, relevant columns like gene identifier, gene symbol, gene description, and GO and KEGG identifiers are automatically extracted. PlantTV also streamlines transcript-to-gene mapping in various ways to produce a gene-centric table, either through GFF or using an additional mapping table. The resulting annotation table can be downloaded and used for downstream analysis.

## Conclusions

We have established PlantTV, a free web-based platform for intuitive exploration of RNA-seq data developed specifically for the plant science community. PlantTV integrates a comprehensive suite of transcriptome analysis tools within a unified interface, enabling dynamic interrogation of transcript abundance plots, differential expression results, volcano plots, heatmaps, functional enrichment analyses, and co-expression networks.

PlantTV addresses a current challenge for working with plant genomes, where for many species multiple annotation files exist and the desired gene information is scattered over multiple tables and resources. To solve these issues, PlantTV contains an integrated annotation builder that allows users to generate and upload custom annotation tables. The annotation builder accommodates varying levels of annotation completeness and accepts multiple input formats, including GFF files, EggNOG and BLASTP outputs, as well as generic tabular data, and allows to add additional gene information from e.g. InterProScan or KAAS through its interactive built-in parsers. For each analysis, users can select their preferred annotation resource.

Moreover, PlantTV was built to promote customized exploration of transcriptomic data, allowing users to easily query the expression patterns of genes, gene families and biological pathways of interest. In this way the platform facilitates hypothesis-driven analyses. In addition to supporting the analysis of private datasets, PlantTV aims to serve as a community resource through the incorporation of publicly available datasets. Once uploaded, these datasets can be explored by all users of the platform, promoting the use and re-use of existing transcriptomic resources within the plant research community.

Overall, PlantTV addresses a critical gap in plant transcriptomics by providing an intuitive, interactive and species-agnostic platform for data analysis. Overall, PlantTV aims to empower plant researchers with and without programming expertise to optimally leverage their RNA-seq data.

## Authors’ contributions

LvB and BDC conceptualized the project. MP and LvB were responsible for software development and manuscript preparation. BDC provided critical revisions to the manuscript. All authors read and approved the final manuscript.

## Acknowledgements

The authors thank Anna Bens and Anke Van Den Berghe for testing PlantTV, Elissar Alam for useful discussions, and Ronny Moreas for deploying PlantTV on the KU Leuven servers.

