## Supplementary Figure 1 for "PlantTV: a versatile user-friendly software to explore RNA-seq data across plant species"

† Corresponding author

**Abstract**

**Background**

Bulk RNA sequencing (RNA-seq) has become a central tool in plant biology for studying genome-wide transcriptional responses. However, interpretation of differential expression results remains challenging, particularly for non-model species where functional annotation resources are fragmented, heterogeneous, or incomplete. Existing user-friendly RNA-seq analysis platforms often rely on predefined annotation databases, limiting their applicability across the diversity of plant species.

**Results**

We developed Plant Transcriptomics Viewer (PlantTV), a free web-based platform for interactive exploration of plant RNA-seq datasets. PlantTV integrates differential expression analysis, expression visualization, functional enrichment analysis, and co-expression network exploration within a single graphical interface. A central feature is the Annotation Builder, which allows users to generate reusable annotation tables from diverse resources, including GFF/GTF files, EggNOG annotations, BLAST results, and custom tabular datasets. Using case studies in *Arabidopsis thaliana*, *Malus domestica*, and *Eucalyptus melliodora*, we demonstrate that PlantTV supports transcriptome analysis across a range of annotation scenarios, from well-annotated reference genomes to de novo transcriptome assemblies.

**Conclusions**

PlantTV provides an interactive and species-independent environment for plant transcriptome analysis. By combining RNA-seq exploration tools with flexible support for custom annotation resources, PlantTV facilitates biological interpretation of transcriptomic datasets in both model and non-model plant species.

**Supplementary Info**

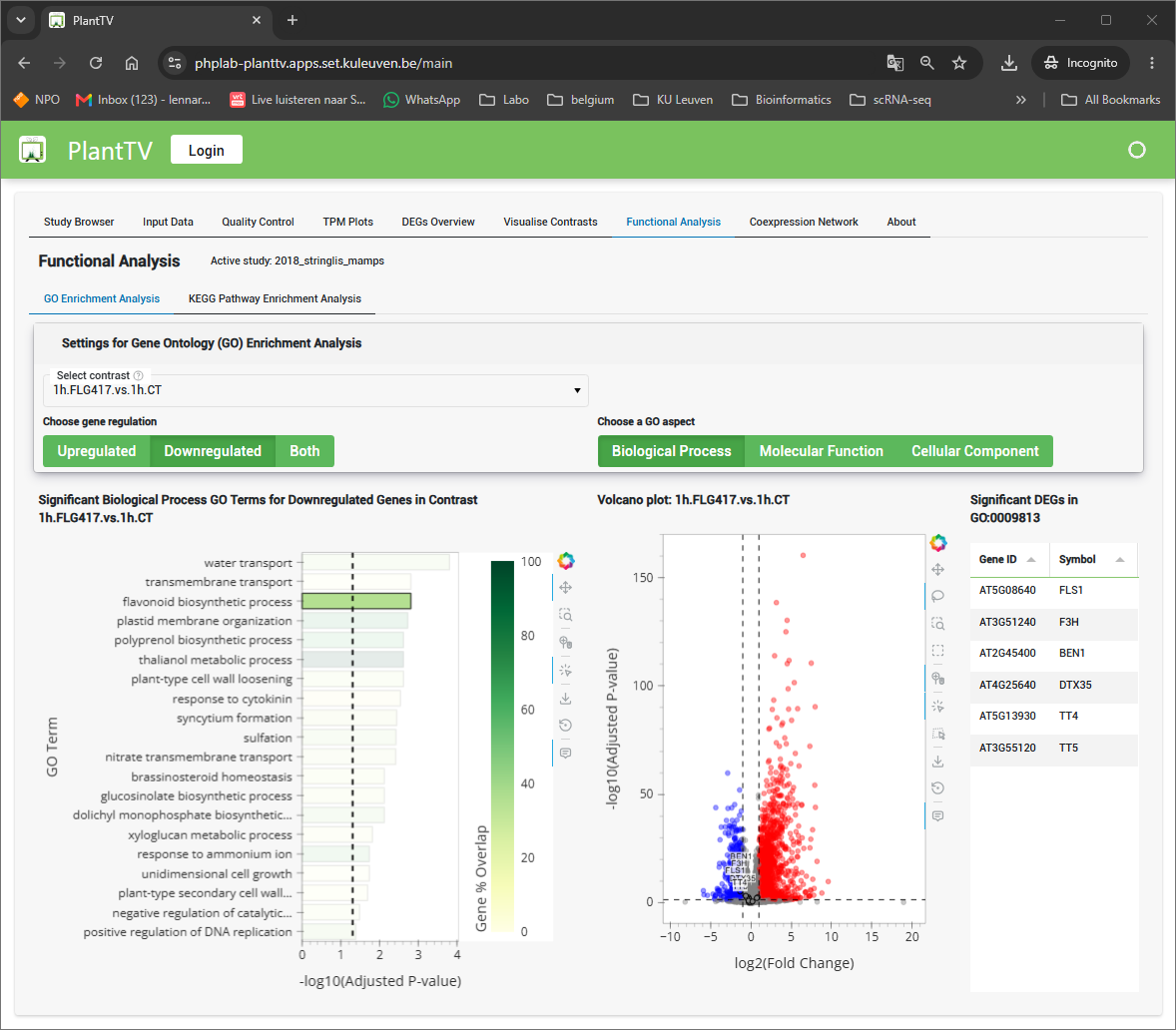

**Fig. S1 | Functional enrichment analysis.** Significantly enriched terms are shown on the left. The GO term Flavonoid biosynthetic process is highlighted and its constituent genes reveal overlap with the cluster found in coexpression network analysis.
